# Harnessing genome prediction in *Brassica napus* through a nested association mapping population

**DOI:** 10.1101/2024.07.29.604859

**Authors:** Sampath Perumal, Erin Higgins, Simarjeet Sra, Yogendra Khedikar, Jessica Moore, Raju Chaudary, Teketel Haile, Kevin Koh, Sally Vail, Stephen J. Robinson, Kyla Horner, Brad Hope, Henry Klein-Gebbinck, David Herrmann, Katy Navabi, Andrew G. Sharpe, Isobel A. P. Parkin

**Affiliations:** Global Institute for Food Security, Saskatoon, Saskatchewan, Canada; Agriculture and Agri-Food Canada, Saskatoon, Saskatchewan, Canada; Department of Plant Breeding and Genetics, Punjab Agricultural University, Ludhiana, India; Nuseed America Inc, Saskatoon, Saskatchewan, Canada; Cargill, Fort Collins, Colorado, USA; Department of Plant Sciences, University of Saskatchewan, Saskatoon, Saskatchewan, Canada; Agriculture and Agri-Food, Beaverlodge Canada, Beaverloge, Alberta, Canada

**Keywords:** Genome prediction, *Brassica napus*, Nested Association Mapping, GWAS, Genomic selection, Canola

## Abstract

Genome prediction (GP) significantly enhances genetic gain by improving selection efficiency and shortening crop breeding cycles. Using a nested association mapping (NAM) population a set of diverse scenarios were assessed to evaluate GP for vital agronomic traits in *B. napus*. GP accuracy was examined by employing different models, marker sets, population sizes, marker densities, and incorporating genome-wide association (GWAS) markers. Eight models, including linear and semi-parametric approaches, were tested. The choice of model minimally impacted GP accuracy across traits. Notably, two models, rrBLUP and RKHS, consistently yielded the highest prediction accuracies. Employing a training population of 1500 lines or more resulted in increased prediction accuracies. Inclusion of single nucleotide absence polymorphism (SNaP) markers significantly improved prediction accuracy, with gains of up to 15%. Utilizing the Brassica 60K Illumina SNP array, our study effectively revealed the genetic potential of the *B. napus* NAM panel. It provided estimates of genomic predictions for crucial agronomic traits through varied prediction scenarios, shedding light on achievable genetic gains. These insights, coupled with marker application, can advance the breeding cycle acceleration in *B. napus*.

**Core ideas:**

- Genome prediction (GP) enhances genetic gains by improving selection efficiency and shortening breeding cycles.
- Factors influencing GP accuracy include model choice, marker types, and population size.
- Inclusion of SNaP markers and highly significant GWAS markers improves prediction accuracy, shedding light on achievable genetic gains.

**Plain Summary:** Genome prediction (GP) is a powerful tool that helps us improve crops more efficiently. In this study, we assessed how well GP works for predicting important traits in *Brassica napus* plants. We tested different models and marker sets to see which ones were most accurate. We found that two models, rrBLUP and RKHS, were consistently the best. Also, including certain types of genetic markers, like SNaP markers and highly significant GWAS markers, improved the predictions. Overall, our study shows that GP can help us understand the genetic potential of *B. napus* plants and improve breeding strategies, which can be exploited to develop better varieties more quickly, which is good news for farmers and the food supply.

## 1 INTRODUCTION

*Brassica napus* (2n=4x=38), commonly known as canola, rapeseed, and oilseed rape, is a crucial oil crop that contributes to global food security. Different ecotypes of *B. napus* exist based on their vernalization requirement: winter type (requiring a long vernalization period), spring type (no vernalization requirement), and semi-winter type (cultivated extensively in China and can flower with or without vernalization) (Hu et al., 2018). Canola provides high-quality edible oil, sustainable biofuel, and protein-rich meal, making it a valuable crop for sustainable food production and efforts are being made to increase its yield, specifically to mitigate the impact of adverse climatic conditions. Understanding the genetic basis of yield and related traits is important for developing high-yielding and climate-smart varieties (Raman et al., 2019). Previous studies have identified important quantitative trait loci (QTLs) associated with grain yield and related traits, such as flowering time and oil content, using genetic mapping in bi-parental populations and diversity panels comprised of cultivars and landraces (Delourme et al., 2018; Fattahi et al., 2018; Pal et al., 2021; Raman et al., 2016; Tian et al., 2019). However, grain yield is a complex trait influenced by multiple small-effect QTLs and genotype-environment interactions. Furthermore, *B. napus*, is an allopolyploid, originating approximately 7500 years ago through hybridization between *B. rapa* and *B. oleracea*, which has resulted in narrow genetic diversity with no known wild populations (Chalhoub et al., 2014). Consequently, because of the complexity of the trait and the low available genetic diversity, achieving substantial genetic gains in grain yield has proven to be challenging.

Advancements in genome sequencing technologies have revolutionized molecular breeding by enabling the development of large numbers of markers at low cost and in less time. The availability of reference genome sequences enabled the design of the Brassica 60K Illumina single nucleotide polymorphism (SNP) array for *B. napus*, which has facilitated the analysis of a large number of samples in a relatively short period of time (Clarke et al., 2016; Mason et al., 2017). This platform has contributed to the identification of QTLs for important traits and the understanding of genetic diversity through genome-wide association studies (GWAS) and genome prediction (GP) (Hurgobin et al., 2018; Qu et al., 2017; Werner et al., 2017).

Additionally, growing evidence suggests that insertion/deletion (Indel) markers, which were previously overlooked due to difficulties in precise identification, contain valuable information for linking traits of interest with the genomic regions controlling their expression (Gabur et al., 2019; Schiessl et al., 2018). Indels and other structural variation impact the detection of bi-allelic markers on SNP arrays such that only one allele is detected, and this has led to a class of marker defined as single nucleotide absence polymorphisms (SNaP) (Gabur et al., 2018). Such markers have been used to identify QTL for agronomically important traits such as disease resistance and yield (Gabur et al., 2018).

Genome-wide association studies (GWAS) are a powerful tool for associating genotypes with phenotypes by leveraging historical recombination events based on linkage disequilibrium (Korte & Farlow, 2013). GWAS allows for the rapid discovery of candidate loci and markers associated with traits of interest, although its resolution depends on the decay of linkage disequilibrium, genetic distance, and marker density (Luo et al., 2019). While GWAS excels in identifying QTLs associated with traits at fine resolution compared to linkage mapping, it is less effective in detecting rare variants (Eichler et al., 2010). However, the identification of rare alleles can be achieved using populations derived from multi-parent crosses, such as Nested Association Mapping (NAM), Multi-parent Advanced Generation Inter-Crosses (MAGIC), and Random-Open-parent Association Mapping (ROAM) populations (McMullen et al., 2009; Sannemann et al., 2015; Xiao et al., 2016). In addition, GWAS has been performed in *B. napus* and multiple other important crops for many different traits, including yield and agronomic traits (Cai et al., 2014; Gajardo et al., 2015; Harper et al., 2012; Hu et al., 2022; Li et al., 2016; Lu et al., 2017; Tian et al., 2019). However, the translation of these findings into practical breeding applications has been limited.

The genetic gain in canola is restricted due to its recent origin, narrow genetic diversity, and selective breeding for specific fatty acid and glucosinolate profiles. NAM populations, developed from multi-parent crosses where founder lines with wide diversity are crossed to a common parent, generate multiple bi-parental recombinant inbred lines (RILs) (Buckler et al., 2009).

NAM populations have the potential to identify traits of interest more effectively than bi-parental populations or diverse lines by leveraging historical and recent recombination events. Additionally, NAM populations are valuable for mapping highly complex and low heritable traits in various crops, including maize, soybean, rice, and *B. napus* (Buckler et al., 2009; Hu et al., 2018; Sharma et al., 2018; Song et al., 2017).

Genome prediction (GP) is a widely used modern breeding approach that uses genotype and phenotype information from a training population to predict genomic estimated breeding values (GEBV) of untested individuals based on genome-wide marker effects (Bhat et al., 2016; Crossa et al., 2017; Heffner et al., 2009). GP has been extensively utilized in animal breeding and offers potential for plant breeding (Desta & Ortiz, 2014; Xu et al., 2020). GP can handle the complexity of plant phenotypes influenced by environmental factors (Bhat et al., 2016). Furthermore, GP helps to accelerate genetic gain by increasing selection gain and selection intensity (Budhlakoti et al., 2022; Voss-Fels et al., 2018; Werner et al., 2018). GP can shorten the breeding cycle by eliminating multiple rounds of phenotype selection using genome-enabled selection (Crossa et al., 2017; Xu et al., 2021). Significant improvements have been achieved in plant breeding, particularly for yield and other agronomic traits, through GP (Bhat et al., 2016; Desta & Ortiz, 2014; Millet et al., 2019). A number of studies have investigated GP for *B. napus* for various agronomically important traits such as yield, disease resistance, and seed quality (Fikere et al., 2020; Fikere et al., 2018; Jan et al., 2016; Roy et al., 2022; Snowdon & Iniguez Luy, 2012; Weber et al., 2023; Werner et al., 2017; Werner et al., 2018; Zou et al., 2016). Various genomic selection methods, including parametric, non-parametric, and machine learning methods, have been developed to assess the prediction accuracy of phenotypes using simulated and empirical data (Azodi et al., 2019; Crossa, Jarquín, et al., 2016; Pérez & de Los Campos, 2014).

In this study, we evaluated the genomic-enabled prediction potential of NAM RILs using markers derived from a 60K Brassica SNP array. We considered various features, including different marker sets, models, marker density, training population size, the inclusion of GWAS results, and the incorporation of SNaP markers, to optimize the genetic potential for yield and yield-related traits.

## 2 MATERIALS AND METHODS

### 2.1 Plant materials

The *B. napus* Nested Association Mapping (SKBnNAM) founder panel of 51 lines was described in Ebersbach et al., (2021). The founders were each crossed to a common parent (NAM 0) resulting in 50 different F_1_ combinations, which through five generations of single seed descent generated the SKBnNAM recombinant inbred lines (RILs) (Figure S1). Approximately 40-60 RILs per family were advanced from the 50 F_1_ combinations, resulting in a final SKBnNAM population comprising 2,572 RILs (Table S1; Figure S1).

### 2.2 Phenotyping

The SKBnNAM population was evaluated for different phenotypic traits at the Agriculture and Agri-Food Canada Llewellyn Farm field station in Saskatoon (latitude 52°04’, longitude 106°10’) in 2017. The field experiment was established in 1.9 m^2^ nursery-plots with a single seeded row with 0.6m spacing, arranged in a Modified Augmented Design type II (You et al. 2013). The whole-plots contained nine sub-plots arranged three by three, with the reference line as a primary check in the center. The whole-plots were arranged as a lattice of 18 rows and 65 columns. Two secondary checks (Founder Line 72 and Founder Line 76) were planted in nine whole-plots distributed across the trial. Four important agronomic traits, including days to flowering (DTF), days to maturity (DTM), plant height (PH), and thousand kernel weight (TKW), were evaluated in this study. DTF was recorded when at least 50% of the plants had a single open flower and the DTM was recorded when seed colour change was evident in pods halfway up the main raceme. Plant height (in cm) was measured as the average the plant height from 3 plants within the plots at physiological maturity. TKW (in grams) was determined by sampling the weight of 1,000 seeds per plot, from the bulk-harvested seed, using an electric balance (Table S1). The phenotypic data of all the NAM RILs were analyzed using an in-house R script using the statistical models previously described (You et al. 2013). Briefly, before proceeding with the genomic analysis, phenotypic data were adjusted for spatial variability within the field, as designated by the modified augmented design. Three methods of adjustment were performed to adjust for spatial heterogeneity. For instance, the first method used the differences between the overall mean of primary checks and the means of primary checks in the row and column corresponding to the main plot position to adjust the entries in the corresponding main plot. The second method involved regressing several secondary checks on the primary checks from the same main plots, calculating the product of this slope and the difference between the primary check of a main plot and the overall primary check mean, and adding the product to each entry in the given main plot. The third method was performed by applying the first method, followed by the second method. Analysis of variance (ANOVA) was performed on the replicated entries in the trial for the original and 3 adjusted data sets. The data set which yielded the smallest error in the ANOVA of replicated entries was used for further analyses (Schaalje et al., 1987; You et al., 2013).

### 2.3 SNP and SNaP Genotyping

The SKBnNAM population was genotyped using the high-density *Brassica* 60K Illumina Infinium™ array, which contains a total of 52,157 SNPs (Clarke et al., 2016; Mason et al., 2017). High-quality DNA was extracted from young leaf tissue using the cetyl tri-methyl ammonium bromide (CTAB) method (Murray & Thompson, 1980), and 200 ng of DNA was hybridized to the Brassica 60K array following the manufacturer’s protocol (Illumina Inc., San Diego, CA). The array was scanned using the Illumina HiScan system, and the SNP data were analyzed using the genotyping module of the GenomeStudio software package (Illumina Inc.) with default settings, except for the no-call threshold, which was set at 0.05. SNP markers with missing data (>5%) and minor allele frequencies (<5%) were removed, resulting in 18,687 markers retained for further analysis. The genome of *B. napus* spring type accession DH12075(v3) was used as a reference (unpublished data; www.cruciferseq.ca). The Mapthin software (www.staff.ncl.ac.uk/richard.howey/mapthin/) was used to reduce the number of markers based on base pair position of the marker, resulting in 4,926 markers for the Genome prediction analysis (Figure 1A, Table S1 and Table S2). The markers were scored as follows: -1 for alternate allele, 0 for heterozygous, and 1 for reference allele.

**Figure 1:**
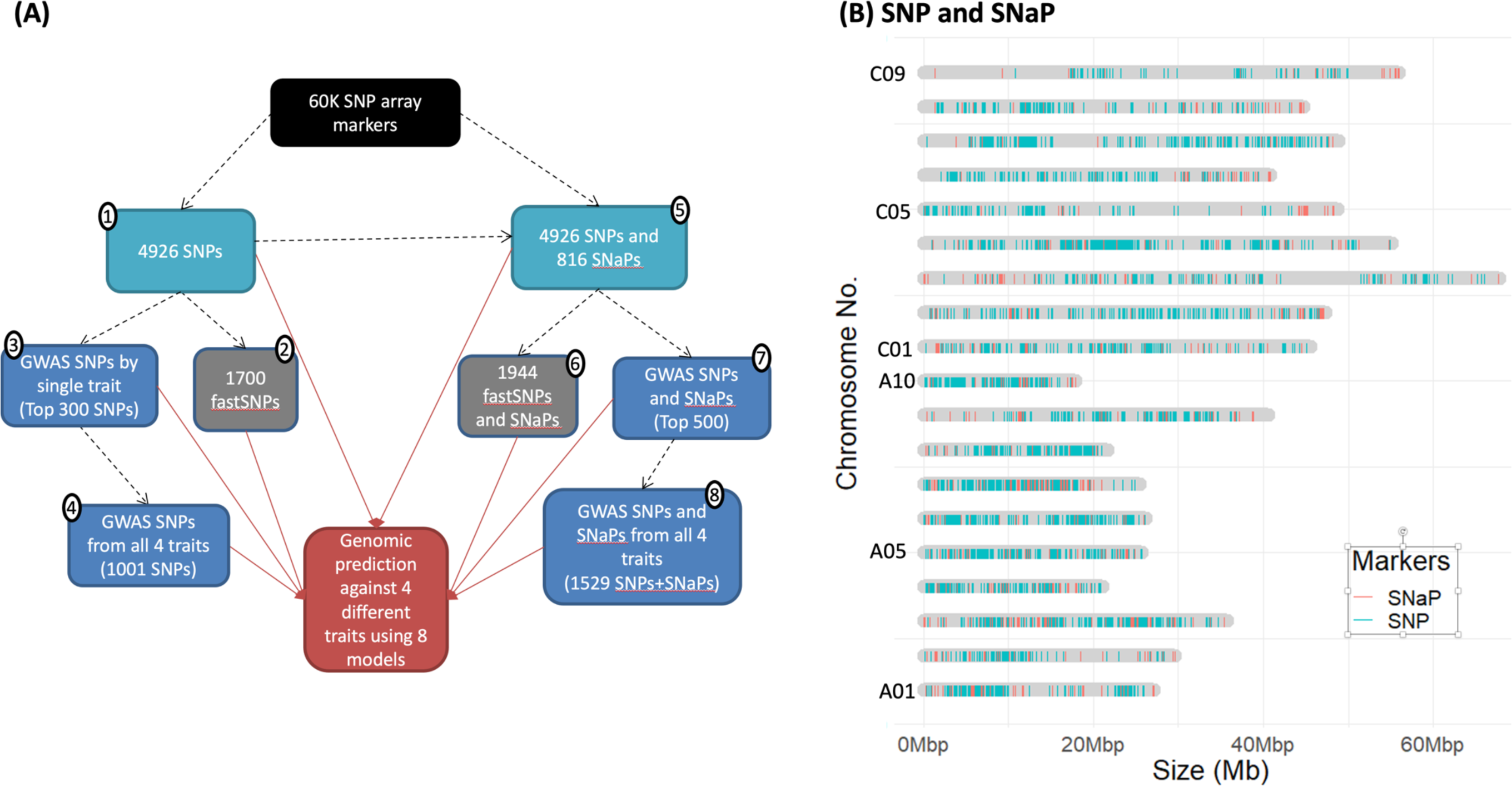
Molecular markers for testing genomic prediction (GP) in Canola. **(A)**Overview of the development of marker datasets for assessing GP accuracy on the *B. napus* NAM population. The dotted line represents the marker development process, and the solid red arrow indicates the utilization of eight different marker sets for the analysis of genomic prediction. Further details on the various marker sets can be found in Table S2. **(B)** Distribution of SNP and SNaP markers along the *B. napus* DH12075 pseudo-chromosomes.

SNaP markers were identified from failed SNP calls of the Brassica 60K Illumina Infinium™ array data, following a previously reported method (Gabur et al., 2018). For example, in a recombinant inbred population, if the marker frequency is 50% for absence (failed calls) and 50% for called alleles, it is considered a SNaP marker. In our analysis, a marker was classified as a SNaP marker if the absence frequency was between 40% and 60% in any one of the 50 families. The absence allele was scored as 1, and the alternate allele was scored as -1.

Furthermore, considering that markers present in genes may have a greater impact on prediction accuracy than markers in non-genic regions, we extracted markers present in genic regions, known as function-associated specific trait (FAST) SNP markers, from the entire SNP and SNaP marker set (Fu et al., 2017). These FAST SNP markers were then used to assess the prediction potential (Figure A1A, Table S1 and Table S2).

### 2.4 Population structure analysis

The genetic diversity of the population was investigated using the fastSTRUCTURE program (Raj et al., 2014). The structure-format SNP+SNaP genotypes were used for STRUCTURE analysis with default parameters. To determine the optimal number of structure model components (K), the analysis was run for 10 replicates (K=1-10). The optimal K value was estimated using the ΔK method implemented in the choose K.py script provided by fastSTRUCTURE. The population structure was visualized based on the optimal K value using the internal fastSTRUCTURE script distruct.py, which utilizes Distruct 2.1 (Rosenberg, 2004).

### 2.5 Genome-wide association studies (GWAS)

Association analyses were performed for each phenotypic trait using both the SNP data alone and the combined SNP and SNaP data (SNP+SNaP). The SUPER algorithm implemented in the Genomic Association and Prediction Integrated Tool (GAPIT) package (Lipka et al., 2012) was used with default parameters to identify SNP effects. The SUPER algorithm increases statistical power by estimating the kinship matrix from associated markers rather than using all markers. Markers showing potential association with phenotypes and thus may have a higher statistical power for providing accurate GP were identified. The markers with the highest effect on the phenotypes, based on low p-value (0.001) were extracted from the GWAS results and utilized in the subsequent GS analysis (Table S2). The top 300 trait-specific GWAS SNPs were extracted for each trait and combined to create the set of all GWAS SNPs (GWAS SNP(all)). Similarly, the top 500 SNP+SNaP markers from GWAS were used to create the set of all GWAS SNP+SNaP markers (GWAS SNP+SNaP(all)) for each trait (Figure 1 and Table S2).

### 2.6 Genome prediction – Models used and methods of evaluation

Several standard single-trait prediction models, including ridge regression best linear unbiased prediction (rrBLUP), genomic best linear unbiased prediction (G-BLUP), BayesA, BayesB, BayesCπ, Bayesian Lasso (BL), Bayesian ridge regression (BRR), and Bayesian reproducing kernel Hilbert spaces regression (RKHS), were used for genomic prediction. All models, except for RKHS, are linear models based on additive genetic effects, while RKHS is a semi-parametric model that accounts for both additive and non-additive genetic effects. The BGLR package v1.0.8 with default parameters (Pérez & de Los Campos, 2014) was used to fit the linear models, except for rrBLUP, which was fitted using the rrBLUP package v4.4 (Endelman, 2011) in the R-environment. All models were run for 30,000 iterations, with the first 10,000 iterations discarded as burn-in. Prediction of lines was performed using 5-fold cross-validation (CV), where the entire population was divided into five mutually exclusive groups, and in each fold the four groups (80% of the lines) were used as the training population, while the remaining group (20% of the lines) was used as the testing population. This was repeated five times so that each group was used as a validation set once. The prediction accuracy of all models was evaluated based on the Pearson’s correlation (r) between the predicted and observed (true) phenotypic values of the lines.

### 2.7 Genome prediction scenarios

Genome prediction accuracies were estimated under five different scenarios, including different marker datasets, population sizes, marker densities, and incorporating population structure**. 1) *Different marker datasets***: The Genome prediction accuracy was estimated using eight different marker datasets: SNP, FAST SNP, GWAS SNP (sp), GWAS SNP (all), SNP+SNaP, FAST SNP+SNaP, GWAS SNP+SNaP (sp), and GWAS SNP+SNaP (all). These marker datasets were used to assess the prediction accuracy for the four different traits (Figure 1A; Table S2). **2) *Marker density*:** The prediction potential was assessed for various marker densities using the SNP+SNaP and GWAS SNP+SNaP datasets. Markers were sub-selected at 25%, 50%, 75%, and 100% density based on inter-marker density. Specifically, for 25%, we systematically selected every fourth SNP. For 50%, we chose every other SNP. The panel for 75% was created by deleting every 4^th^ SNP, and for 100%, the entire set of SNPs was included. **3) *Size of the training population*:** The optimal training population size was evaluated by analyzing subsets of 500, 1000, 1500, and 2000 lines as the training population, with the remaining lines serving as the validation population. The training population included lines representing all 50 RIL families. **4) *Size of the population***: To determine the optimal population size for achieving better prediction accuracy, subsets of 500, 1000, 1500, 2000, and >2000 lines were tested using the eight different models and the GWAS SNP+SNaP and SNP+SNaP datasets with 5-fold cross-validation.

## 3 RESULTS

### 3.1 Extraction of high-quality SNP+SNaP markers for the *Brassica napus* NAM panel

The SKBnNAM population consists of 2,572 lines, which were used to evaluate the Genome prediction potential for four traits using eight different models under different scenarios. After thinning markers extracted from the Brassica 60K Illumina Infinium chip analyses resulted in 4,926 high-quality SNP markers (Figure 1A and Table S2). Additionally, 816 SNaP markers were identified based on segregating null (failed call) alleles with a frequency threshold for failed calls between 40-60% in any one RIL sub-population. SNP and SNaP markers were distributed across the whole genome, with an average of 299 markers per chromosome and a density of approximately 6 markers per Mb. Chromosome BnaC4 had the highest number of markers (704), while chromosome BnaA2 had the lowest (140). Interestingly, SNaP markers were observed in regions where SNPs were limited or absent (Figure 1B). The average polymorphic information content was 0.34 for SNP markers and 0.32 for SNaP markers.

### 3.2 Population structure of *Brassica napus* NAM population

Population structure analysis of the SKBnNAM panel revealed two distinct groups and a third admixture group (Figure S3A). The optimal K analysis indicated that a model complexity that maximizes marginal likelihood occurs when K = 2, suggesting that two clusters most likely explain the population structure of the BnNAM panel. The maximum population structure could be identified when the model components were set as K = 9 (Figure S3B). Principal component analysis (PCA) showed a similar pattern to the population structure analysis, with two major groups and a minor group (Figure 2). Group 1 contained the majority of lines (2,122) and contributed over 50% of the genetic diversity, followed by group 2 with 426 lines, contributing approximately 49% of the genetic diversity. Linkage disequilibrium (LD) analysis revealed that LD dropped to half of its maximum value at a distance of approximately 550 kb (Figure S4).

**Figure 2:**
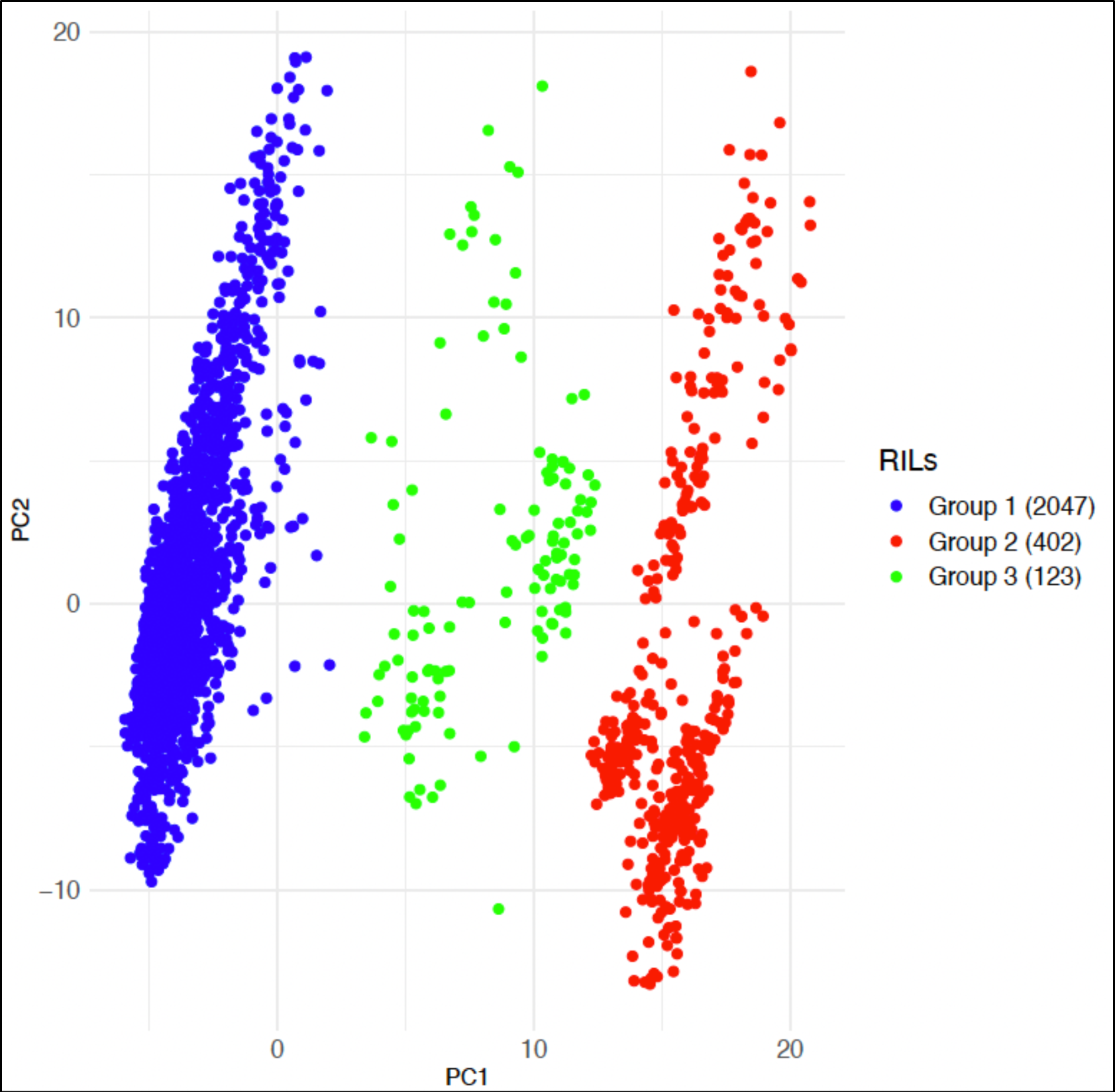
Principal component analysis of 2623 NAM RILs based on 5743 SNP and SNaP markers.

### 3.3 Genome-wide association analysis (GWAS)

Among four significant agronomic traits focused on this study, DTF was a highly heritable trait (h^2^=0.74), followed by PH (0.61), TKW (0.58), and DTM (0.43). There was wide variation for plant height, ranging from 91 to 200 cm (Figure 3). DTF showed a positive correlation with DTM, but no significant correlation was observed with PH or TKW. Additionally, an increase in TKW was observed with an increase in plant height.

**Figure 3:**
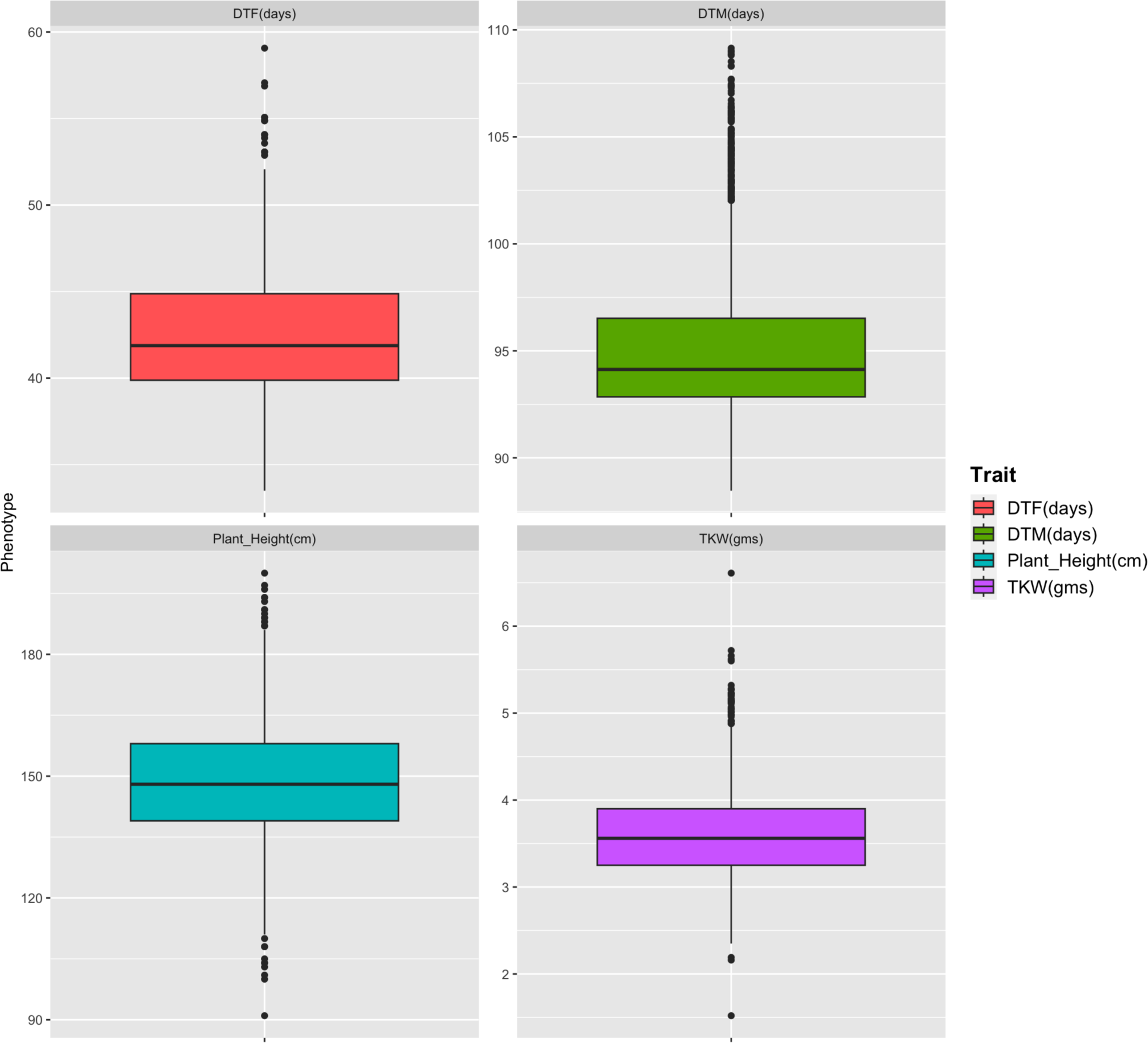
Distribution of phenotypes for four different traits utilized in this study. The boxplot illustrates quartile values for each phenotype, with whiskers indicating the approximate 95% confidence interval. The median is marked by the mid-box, while the mean value is represented by the horizontal line within the mid-box. Outliers are indicated by dots. Traits: DTF (days to 50% flowering), DTM (days to maturity), Plant height (in cm), TKW (Thousand kernel weight in grams).

GWAS was performed for the four different traits using two different marker sets: SNP and SNP+SNaP (Figure 4). GWAS was conducted to understand the impact of SNaP markers on trait association compared to SNP markers alone. Based on the 4,926 SNP marker set, three regions were significantly associated with the four phenotypes. Particularly, a region on chromosome 2 (SNP: Bna.A02.p1585968) was significantly associated with three different traits: DTF, DTM, and PH. The SNP was present in the gene BnaA02g003060.1DH, which is a homolog of the RPN5A gene that functions in ATP-dependent degradation of ubiquitinated proteins (Table S3). Similarly, GWAS using the SNP+SNaP marker set, which contained 5,742 markers (816 from SNaP), revealed many additional regions associated with the four traits. One additional region on chromosome Bna.C09 was significantly associated with the DTM trait, and two other regions on chromosome Bna.C09 were associated with DTF and PH traits (Figure 4B). These regions were not detected by SNP markers alone, suggesting the importance of SNaP markers. Furthermore, functional annotation of the significant markers associated with traits showed that the genes were related to various plant growth, transcription regulation, and metabolism processes (Table S3).

**Figure 4:**
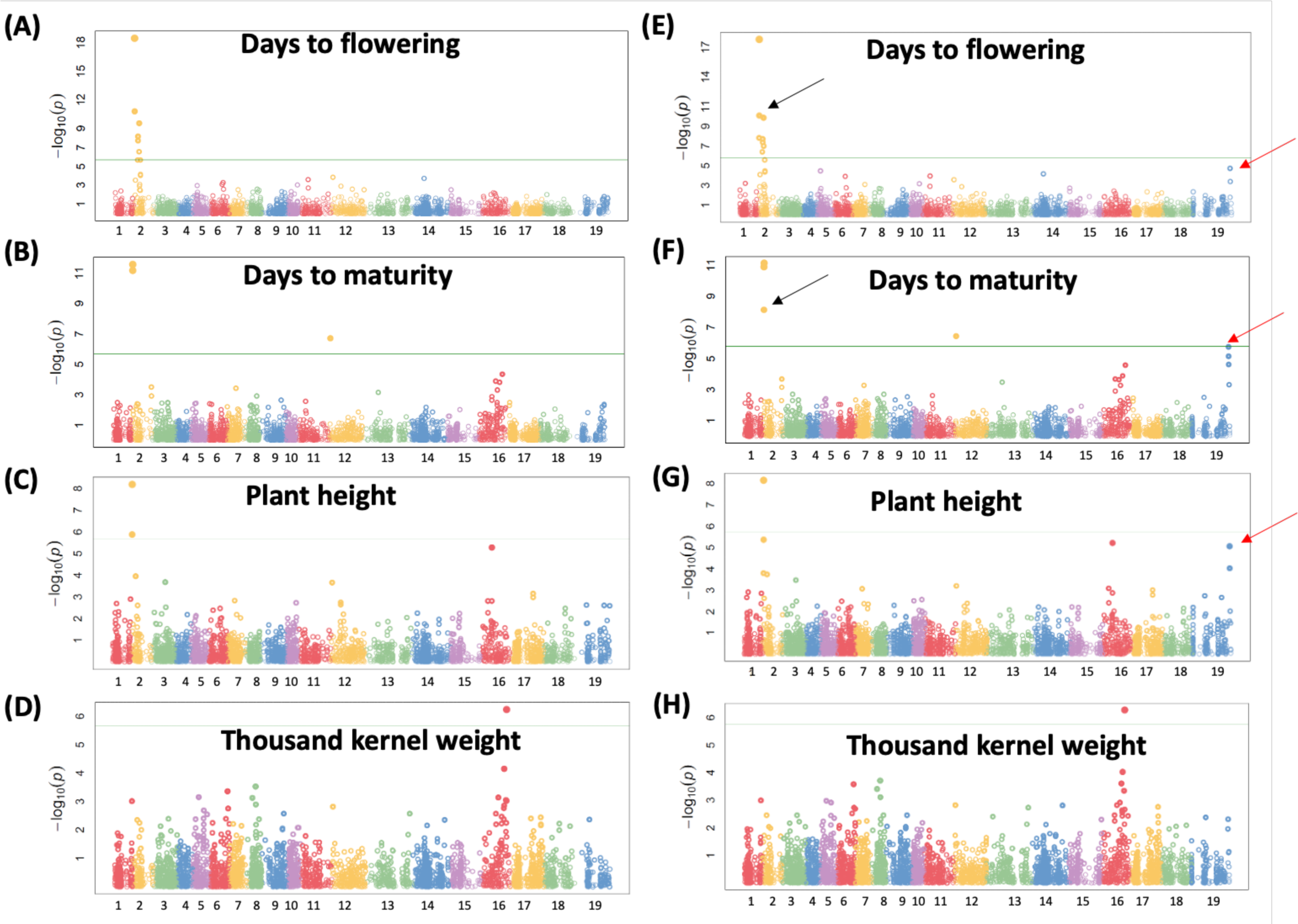
Genome-wide association studies for four different traits using SNP (A-D) and SNP+SNaP (E-H) markers. The green horizontal line indicates the significance threshold line based on the False discovery rate (FDR) of 5, markers above the line are significantly associated with the indicated trait. Black arrows indicate the addition of significant SNaP markers, red arrows indicate regions only identified by SNaP markers. X-axis shows the physical position of the markers on the chromosomes: 1-10 -BnaA01-A10 and 11-19 - BnaC01-C9.

### 3.4 Genome prediction of SKBnNAM panel with eight different models

The overall prediction accuracy for each trait ranged from moderate to high (0.27-0.71) across the different models used (Figure 5; Table S3). Among the four traits, DTF had the highest prediction accuracy (0.71), followed by plant height (0.63). TKW showed the lowest prediction accuracy, ranging from 0.32 to 0.40, followed by DTM with 0.36-0.46 (Figure 5; Table S3).

**Figure 5:**
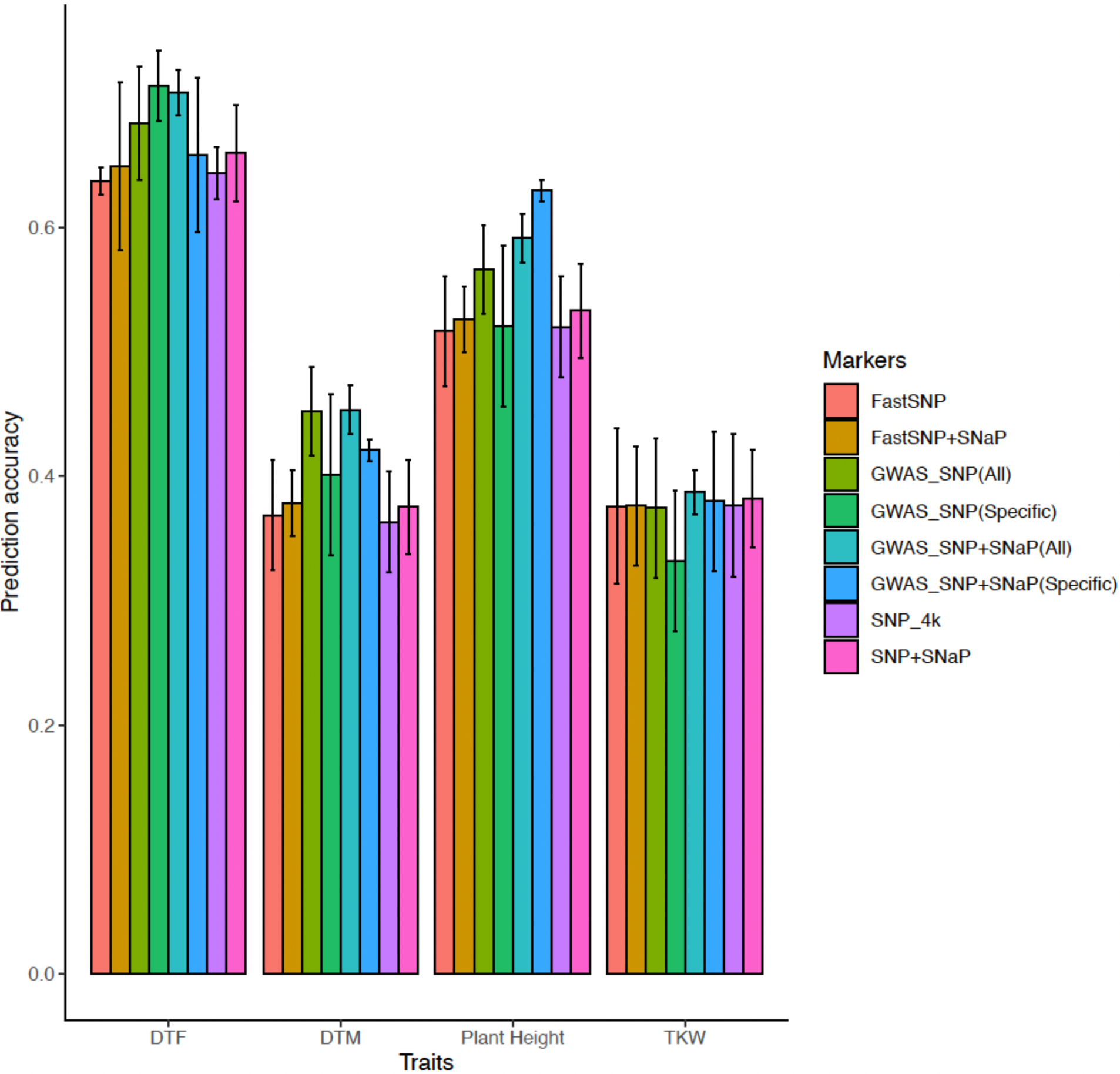
Genome prediction accuracy of *B. napus* NAM panel on eight different marker sets using rrBLUP.

Traits with high heritability, such as DTF and PH, had higher prediction accuracy compared to low heritability traits. There were no significant differences in prediction accuracy among the eight different models used. In most cases, the differences were 2-4% between them (Figure 5B). However, the RKHS model consistently produced the highest accuracy for almost all the traits and different marker sets used in the genomic prediction, followed by rrBLUP.

#### 3.4.1 Impact of marker types on prediction accuracy

The prediction potential for the SKBnNAM lines for the four traits was tested based on eight different marker datasets, including SNP, FAST SNP, GWAS SNP (sp), GWAS SNP (all), SNP+SNaP, FAST SNP+SNaP, GWAS SNP+SNaP (sp), and GWAS SNP+SNaP (all) (Figure S2; Figure 5; Table S3). The total number of markers varied from 300 to 5,742 across the eight different marker sets. GWAS SNP+SNaP markers showed a similar distribution across the genome as the originally selected SNP+SNaP markers, with an average of 1.4 markers per Mb (Figure S2). The 1,944 markers (34%) identified as FAST SNP markers, which were present in genes, were mostly found in the intronic regions (1,098 (57%)) rather than exons of the annotated genes (Table S2).

Different marker sets provided varying levels of prediction accuracy for each trait. Among the eight different marker sets used, the SNP combined with SNaP marker set (SNP+SNaP) yielded the highest prediction accuracy compared to SNP-only marker datasets (Figure 5; Table S3). We found differences between 2 and 10% when using SNP+SNaP markers for plant height (0.53 vs. 0.63). The GWAS SNP+SNaP (all) marker set produced the highest prediction accuracy (0.71-0.72) for DTF, which was similar to the 300 GWAS trait-specific SNPs (GWAS SNP (Sp)) (0.71), while other datasets showed similar or slightly lower prediction accuracy (0.64-0.70). For DTM, the GWAS SNP+SNaP (all) marker set predicted the highest accuracy (0.46), which was also comparable to GWAS SNP (all) (0.44-0.45). Interestingly, for PH, the highest prediction accuracy was observed with the GWAS SNP+SNaP (Sp) marker set compared to all other different marker datasets. This suggests that the capture of significantly associated presence/absence alleles (SNaP markers) improved the prediction accuracy for this trait. For TKW, the prediction accuracy ranged from 0.27 to 0.40, and except for GWAS SNP, every marker set predicted at least 0.37 accuracy for one or more models. Overall, SNP and SNP+SNaP markers had similar prediction accuracies, while markers distributed in the genic regions (FastSNP/FastSNP+SNaP) showed slightly lower prediction accuracy compared to GWAS-based markers, but comparable prediction accuracy to SNP and SNP+SNaP markers (Figure 5; Table S3). Marker sets based on GWAS SNP+SNaP yielded the highest prediction accuracy, except for the plant height trait.

#### 3.4.2 Impact of marker density on prediction accuracy

The prediction potential did not correlate with an increase in marker numbers. For example, the SNP+SNaP marker set with 5,742 markers showed very similar or slightly lower prediction accuracy compared to the GWAS-based marker sets with 300-1,529 markers. Furthermore, even with just 500 markers from GWAS alone, reasonable prediction accuracy was consistently achieved. In fact, an increase in marker density might negatively influence the prediction potential. Testing the prediction potential of the GWAS SNP+SNaP (all) marker set with different subsets of markers demonstrated that approximately 50% (765 markers) of the markers were sufficient to achieve reasonable prediction accuracy, while increasing the number of markers beyond 75% (1,147 markers) resulted in decreased or no change in prediction accuracy for all traits except PH (Figure 6). PH showed a sudden increase in accuracy (0.4-0.65) with added markers, suggesting a significant influence of some loci for this particular trait. Similar trends were found for DTF when using the SNP+SNaP marker set (Figure S6).

**Figure 6:**
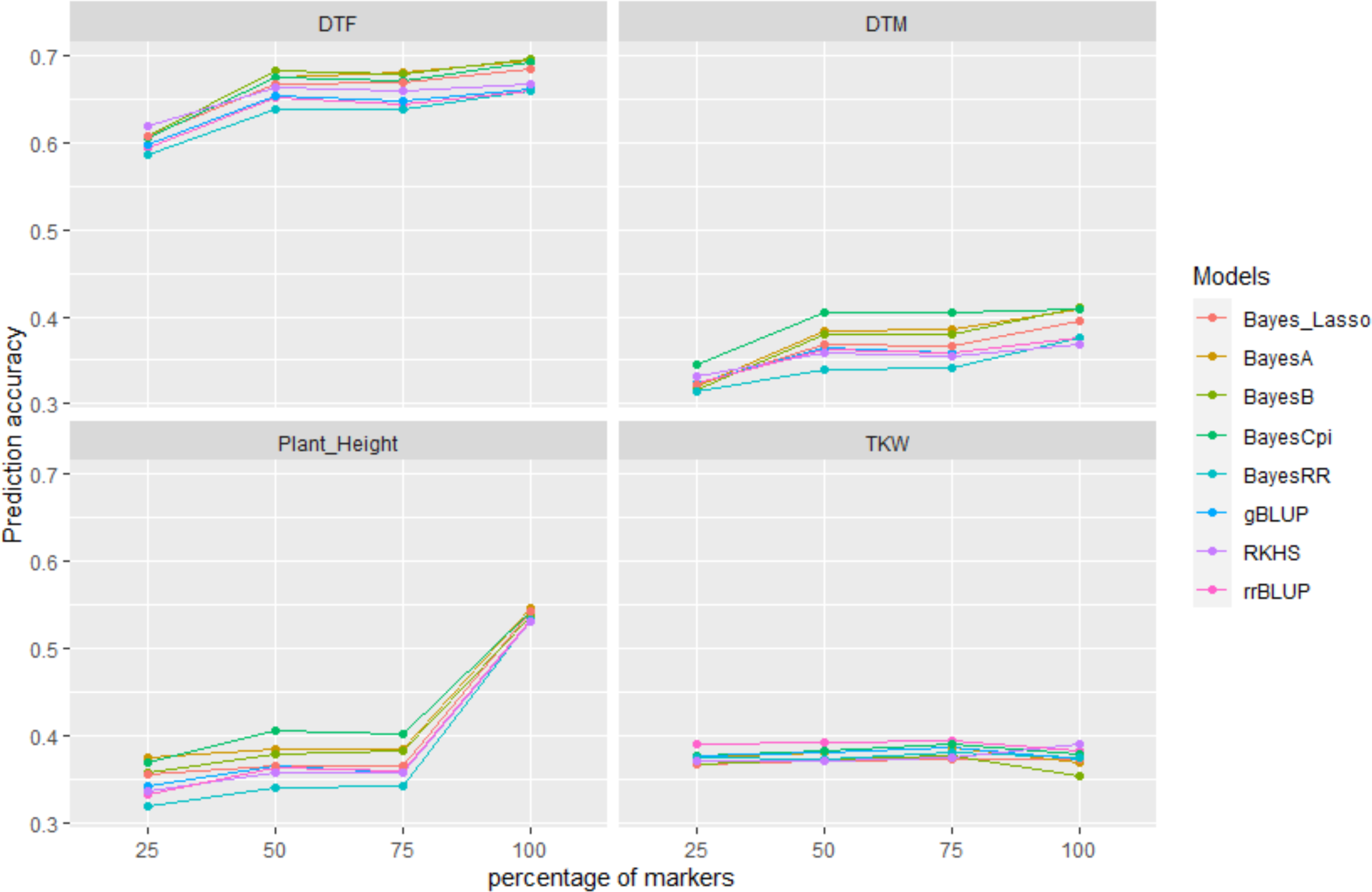
Marker density and genomic prediction accuracy evaluation in *B. napus* NAM panel. Prediction accuracy of GWAS SNP+SNaP marker set was assessed across four different combinations (25% - 100%) for the four traits.

#### 3.4.3 Impact of training population size on prediction accuracy

Prediction accuracy increased as the training population size increased for the GWAS SNP+SNaP marker set across all four traits and eight different models (Figure 7). An initial training population size of 500 was selected, when this was doubled (1,000 lines), the prediction accuracy increased by an average of 5-15%, while tripling the training population (1,500 lines) or increasing by fourfold (2,000 lines) further increased the accuracy by 5-10%. A similar positive trend was observed for the SNP+SNaP marker set (Figure S7). Both marker sets indicated that a training population size between 1,500 and 2,000 lines, representing a minimum ratio of 3:2 (ie.500:1000 lines) for training to test population might be needed to achieve the best prediction accuracy.

**Figure 7:**
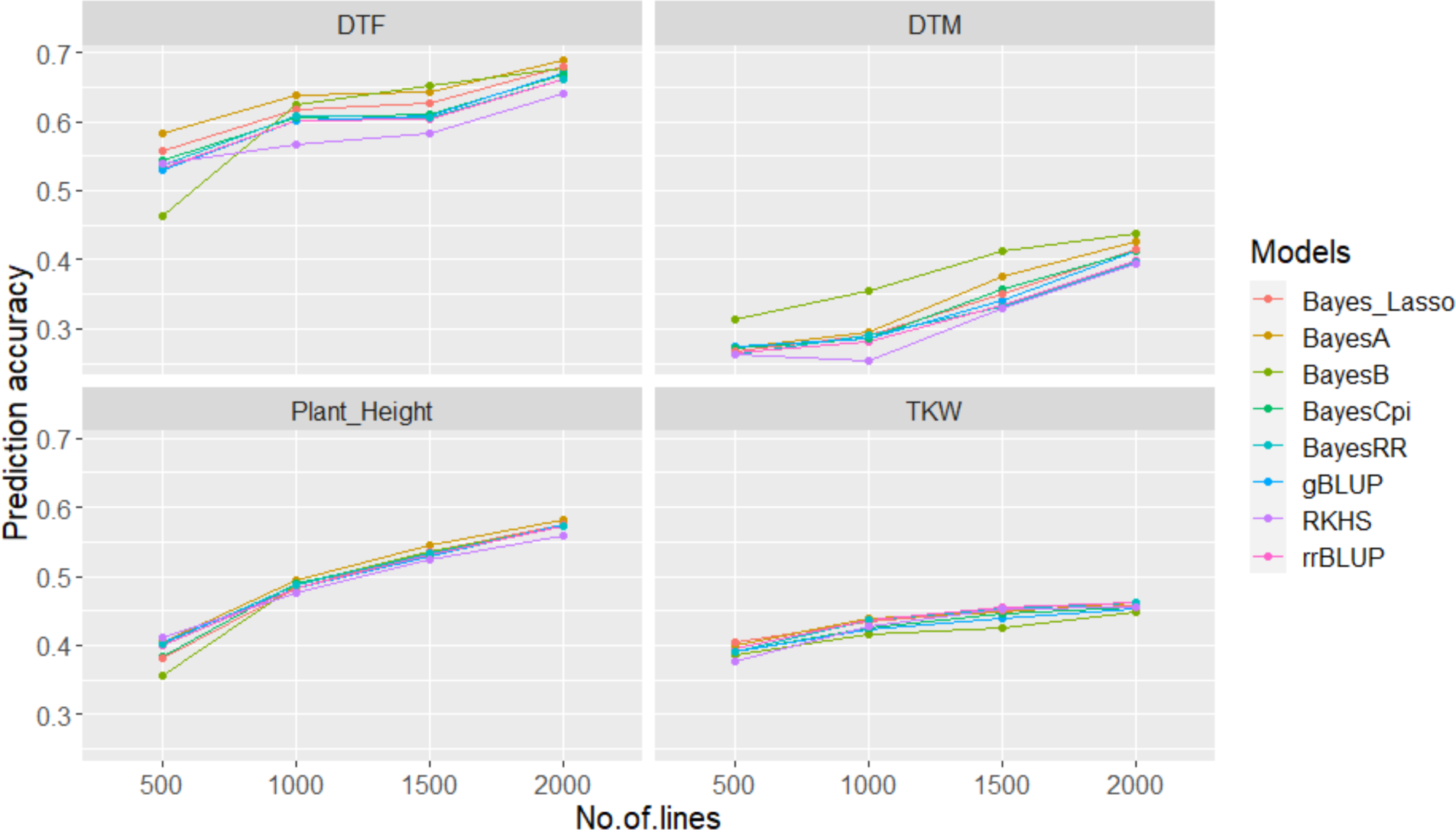
Impact of training population size on prediction accuracy using GWAS SNP+SNaP Marker Set. Prediction accuracy was evaluated for varying sets of lines (500, 1000, 1500, and 2000) to estimate the accuracy on 500 randomly selected testing lines, employing 8 different models.

#### 3.4.4 Impact of absolute population size on prediction accuracy

Prediction accuracies based on different numbers of lines (250 to 2,500) but the same GWAS SNP+SNaP (all) marker set were tested; in each instance 80% of the subset population was used as the training population (Figure 8). In general, prediction accuracy increased as the number of lines increased, and a minimum of 1,000 lines produced reasonable prediction accuracy regardless of the genomic prediction model used (Figure 8).

**Figure 8:**
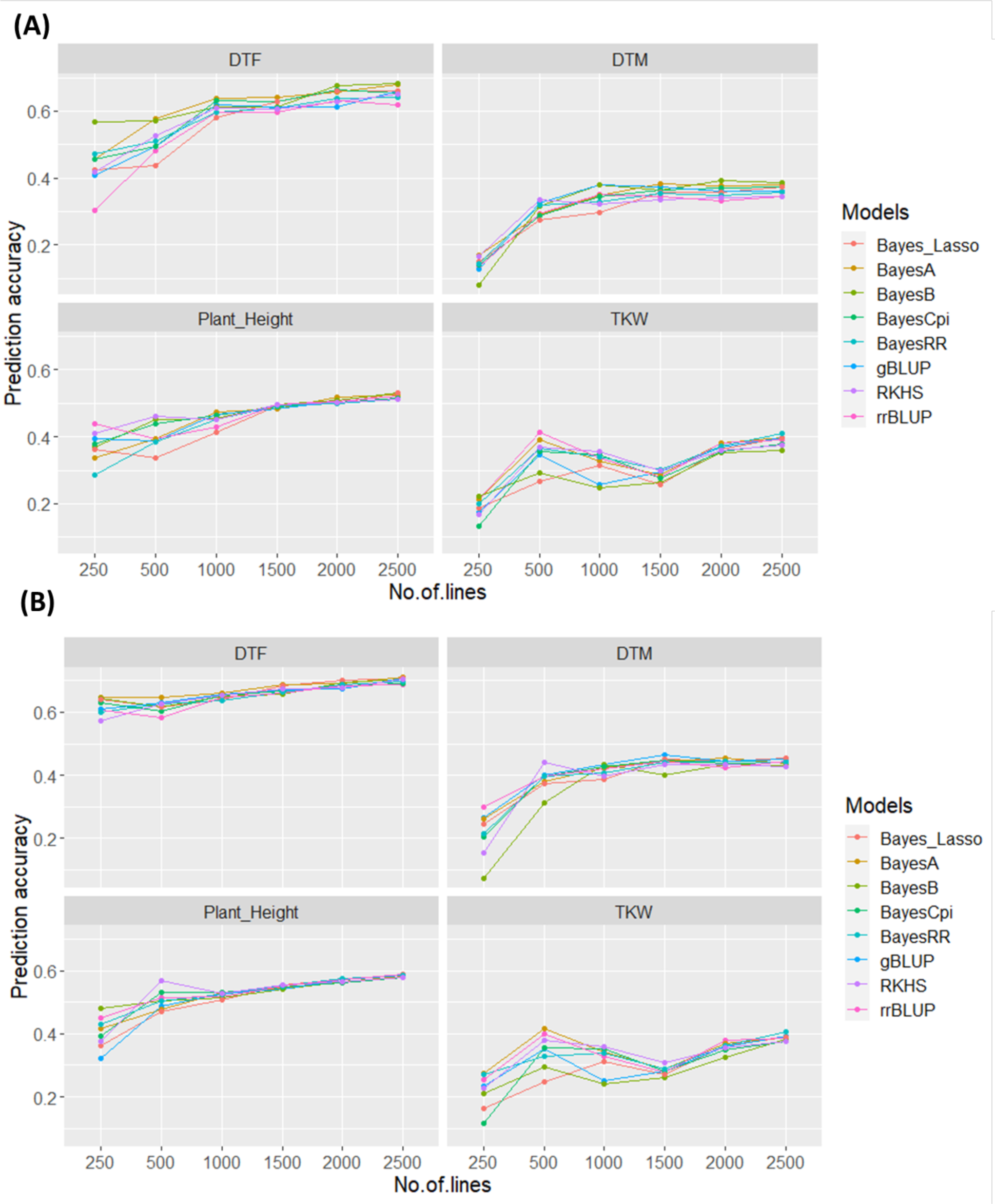
Estimating population size on prediction accuracy based on all SNP+SNaP marker set (A) and GWAS SNP+SNaP marker set (B) using 8 different models. Different set of lines (250, 500, 1000, 1500, 2000 and 2500) were used to estimate the prediction accuracy. In each combination, 80% were used as training set to estimate the prediction accuracy on the remaining 20% of the lines.

## 4 DISCUSSION

Genome prediction offers a significant opportunity to enhance crops by improving selection efficiency and reducing the breeding cycle (Crossa et al., 2017; Rajkumar, 2022; Sharma et al., 2022). Furthermore, GP is a more reliable method and outperforms conventional breeding approaches such as pedigree and QTL-based approaches (Muir, 2007; Nakaya & Isobe, 2012). In addition, GS helps to incorporate unphenotyped and novel germplasm into a breeding program helping to increase the genetic gain and diversity (Budhlakoti et al., 2022). *B. napus* has a narrow genetic diversity, and a few studies have explored its prediction potential using GS (Li et al., 2016; Rajkumar, 2022; Werner et al., 2017; Werner et al., 2018). RILs developed from NAM populations have broad-genetic diversity due to the intensive reshuffling of the genome making them interesting tools to test the performance of genomic-enabled prediction. Although a few studies have explored the impact of marker density, population size, and different models for genomic prediction in canola (Fikere et al., 2020; Weber et al., 2023), prediction potential has not been comprehensively evaluated in *B. napus*, especially using SNaP markers, adjusting marker density and population size, and utilizing some of the more recently adopted models. In this study, genomic prediction potential was comprehensively evaluated using a *B. napus* NAM population for four different traits using eight different marker sets with eight models, including linear and semi-parametric models. Overall, moderate to high prediction accuracies were observed with DTF showing the highest prediction accuracy of 0.72 among the four traits analyzed. Compared to previously available predictions for *B. napus* (Jan et al., 2016; Werner et al., 2018), these analyses showed higher prediction accuracies for DTF (0.56 vs 0.71) and plant height (0.5 vs 0.63). TKW showed the lowest prediction accuracy among the four traits analyzed (0.40). This is attributed to its highly quantitative nature, corroborated by the GWAS analysis (lowest p-value of 5.39E-07). Environmental factors, multigene and low heritable traits can influence the prediction accuracy (de Los Campos et al., 2015). Hence, incorporation of multiple environments in the prediction model might help to increase the prediction accuracy of highly quantitative traits such as TKW (Crossa, de los Campos, et al., 2016; de Los Campos et al., 2015; Haile et al., 2020).

Several factors can influence prediction accuracy, such as the selection of prediction models, marker type and density, relationship between training and testing populations, size of the training population, population structure, linkage disequilibrium, and genotype-by-environment interactions (Crossa et al., 2017; de Los Campos et al., 2015; Luo et al., 2017). A number of these factors were tested utilizing the available SKBnNAM data. The eight different models employed had a modest influence on prediction accuracy. However, among the eight, RKHS demonstrated a higher and more consistent prediction accuracy than the other seven models used. In addition, rrBLUP also showed quite similar prediction accuracy compared to RKHS. Based on our findings we found that RKHS and rrBLUP are the best-performing models for *B. napus*; however, testing of models which possess multiple-trait prediction (Gianola & Fernando, 2020) and machine learning (Azodi et al., 2019; Wang et al., 2020) might help to further improve prediction accuracy.

The relatedness and size of the population can significantly impact GP analysis (Habier et al., 2007; Nakaya & Isobe, 2012). Sufficient marker diversity and a suitably sized training population are important to capture the entire population structure, which can then be tested on random population sets. Using multiple training sets suggested that a minimum of 500 lines is sufficient to achieve reasonable prediction accuracy, and a training population size of 1,000 seemed to be optimal for the *B. napus* population used in these analyses (Figure 6). However, when the number of lines exceeded 2,000, the prediction potential started to decline for two of the four analyzed traits, indicating increased complexity of trait and marker associations (Xavier et al., 2016). It seems essential to consider scaling the training population size linearly with marker density for optimal advantage. Similarly, the analysis of absolute population size revealed an increasing trend as the size of the population increased, with a population of 1,500 or more being beneficial for prediction (Figure 7).

It has been suggested that an increase in the number of markers does not always improve prediction potential, since it can increase the statistical complexity, which may negatively impact prediction accuracy (VanRaden et al., 2011). On the other hand, prediction based on markers linked to a phenotypic trait can improve prediction accuracy. The prediction accuracy also depends on the major or minor effect of the markers (O’Connell et al., 2022). The prediction accuracy started to decline, particularly for the DTF and PH traits, when markers exceeded 1,500; however, an increase in prediction accuracy was observed for TKW as the number of markers increased (Figure 8). This suggests that TKW is controlled by many minor effect QTLs that were more favourably represented as the marker coverage increased, contributing to the improved prediction accuracy.

It is important to select the right combination of markers to achieve better prediction accuracy. Various methods have been employed to reduce the number of markers, which helps to improve computational efficiency. For example, the number of markers can be reduced by selecting a random set from a pool. Additionally, selecting the desired number of markers based on the size of the genome (i.e., 5 markers/Mb region) can be beneficial. However, randomly filtering out markers does not always yield desirable prediction accuracy. Selecting markers based on their effect on traits is crucial and can improve prediction accuracy (Fu et al., 2017; Werner et al., 2018). By selecting markers based on their association with phenotypic traits, in this instance highly significant makers from GWAS analysis, provided an opportunity to eliminate markers with limited potential and reduce the statistical complexity of prediction. As a result, an increased prediction accuracy was observed even with the lowest number of markers (500 markers) for DTF (0.72) (Figure 5; Table S4). This approach will help to reduce the number of markers required for GS, ultimately reducing cost and time. However, it is important to exercise caution when selecting the number of markers, as the elimination of markers potentially associated with minor effects could occur.

Structural variants or indel markers have been largely ignored due to the complexity of their identification using short read sequencing platforms. However, recent studies suggested such markers can have a significant effect and help identify traits of interest (Gabur et al., 2018; Gabur et al., 2019). Recent analyses have shown that SNaP markers derived from SNP arrays can identify QTLs associated with disease resistance in *B. napus* (Gabur et al., 2018).

Incorporating SNaP markers in GWAS analyses helped to identify markers significantly associated with the PH trait which otherwise might have been overlooked. Moreover, a few additional SNaP markers were highly correlated with regions previously identified by SNPs for the DTF and DTM traits. Furthermore, SNaP markers were included in the prediction analysis, resulting in increased accuracies of up to 15% compared to using SNP markers alone. This strongly supports the notion that SNaP markers can greatly influence genome prediction and trait identification particularly in regions where SNP markers were scarce, thus amplifying genetic gains in the study (Weber et al., 2023).

## 5 CONCLUSION

GP has the potential to increase selection efficiency, genetic gain, and reduce the breeding cycle time. This study included a comprehensive analysis of the GP potential for four traits in a *B. napus* NAM population using different types of markers developed from the *Brassica* Illumina 60K SNP array. Further, the prediction potential was evaluated under various scenarios, including different prediction models, population density, marker density, and marker combinations, including SNaP markers. This study demonstrated the impact of SNaP markers on prediction accuracy, resulting in possible increases of up to 15%, especially harnessing the genetic gains overlooked by SNP markers. This finding opens avenues for future research exploiting the potential of SNaP markers or other structural variation markers. Additionally, it is likely that incorporating environmental interactions and exploring alternative prediction models, such as machine learning techniques, could further improve prediction accuracy.

## AUTHOR CONTRIBUTIONS

S.P and I.A.P played roles in designing the project and objectives. E.H and S.S generated the SNP data. S.V designed and implemented the field trial, coordinated collection of the phenotype data and subsequent data analyses, while Y.K and H.K.G curated the phenotype and genotype dataset. I.A.P, S.V, S.J.R, K.H, and B.H developed the NAM population. S.P led the bioinformatic analyses and drafted the manuscript. T.H, K.K and R.C helped with bioinformatic analysis and data visualization. K.N, J.M, and D.H lent their expertise to the genomic prediction analyses. A.G.S, I.A.P and S.P edited the final draft of the manuscript.

## ACKNOWLEDGEMENTS

Sampath Perumal acknowledges the MITACS elevate fellowship supported by MITACS and Cargill Canada. This work was supported by funding from the MITACS and Cargill Canada. The development of the NAM population and all associated data was supported by a grant from the Agricultural Development Fund #20110155 from the Ministry of Agriculture, Saskatchewan, along with additional funding from SaskCanola, and four industry partners.

## SUPPORTING INFORMATION

**Figure S1:**
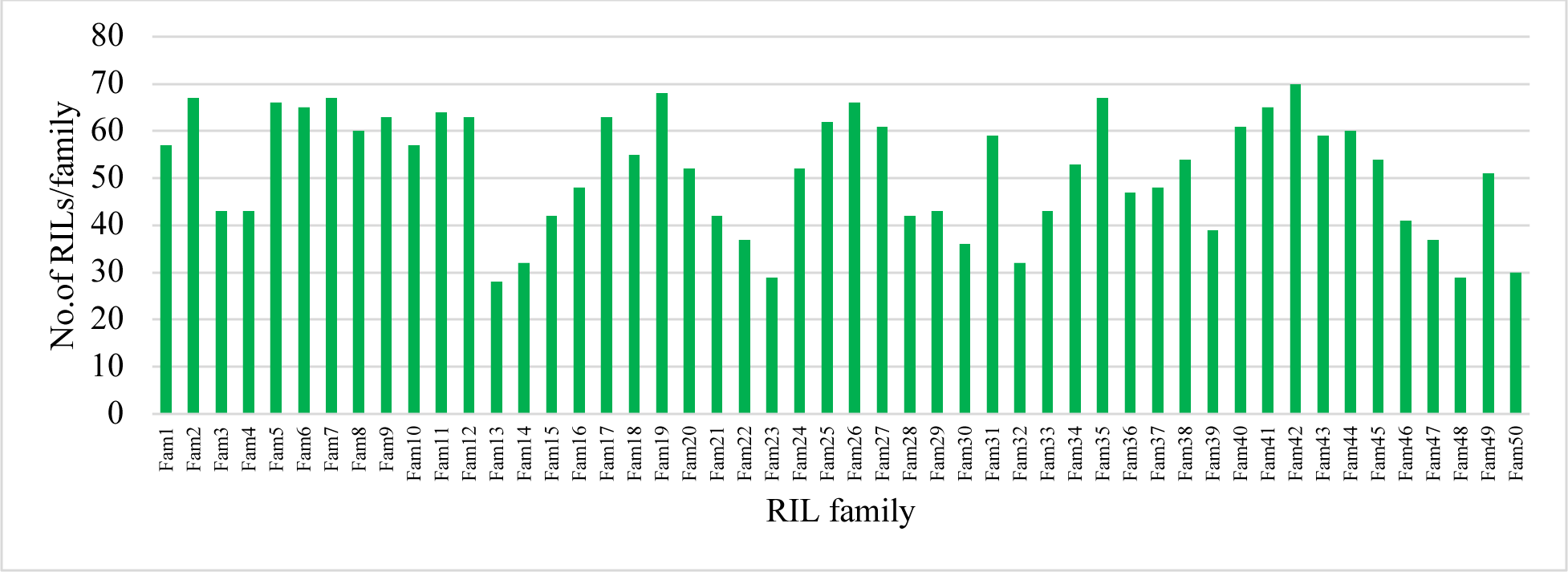
Distribution of number of recombinant inbred lines (RILs) in each *B. napus* NAM family.

**Figure S2:**
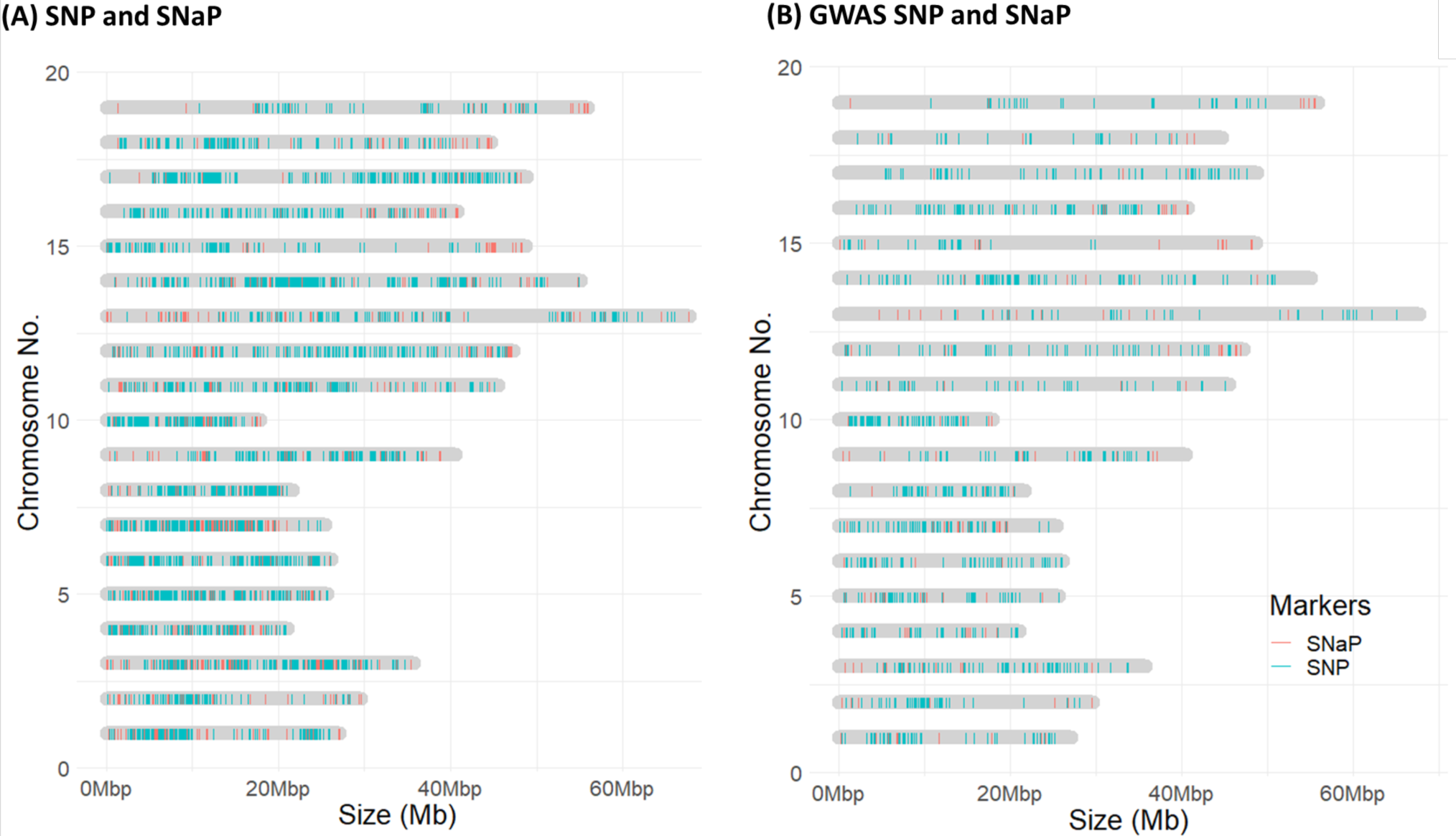
Distribution and comparison of various markers, (A) Total SNP and SNaP (B) GWAS based selection of SNP and SNaPs on the pseudo-chromosome of *B. napus* reference genome (DH12075 v3). For additional details on different marker sets, please refer to Table S2.

**Figure S3:**
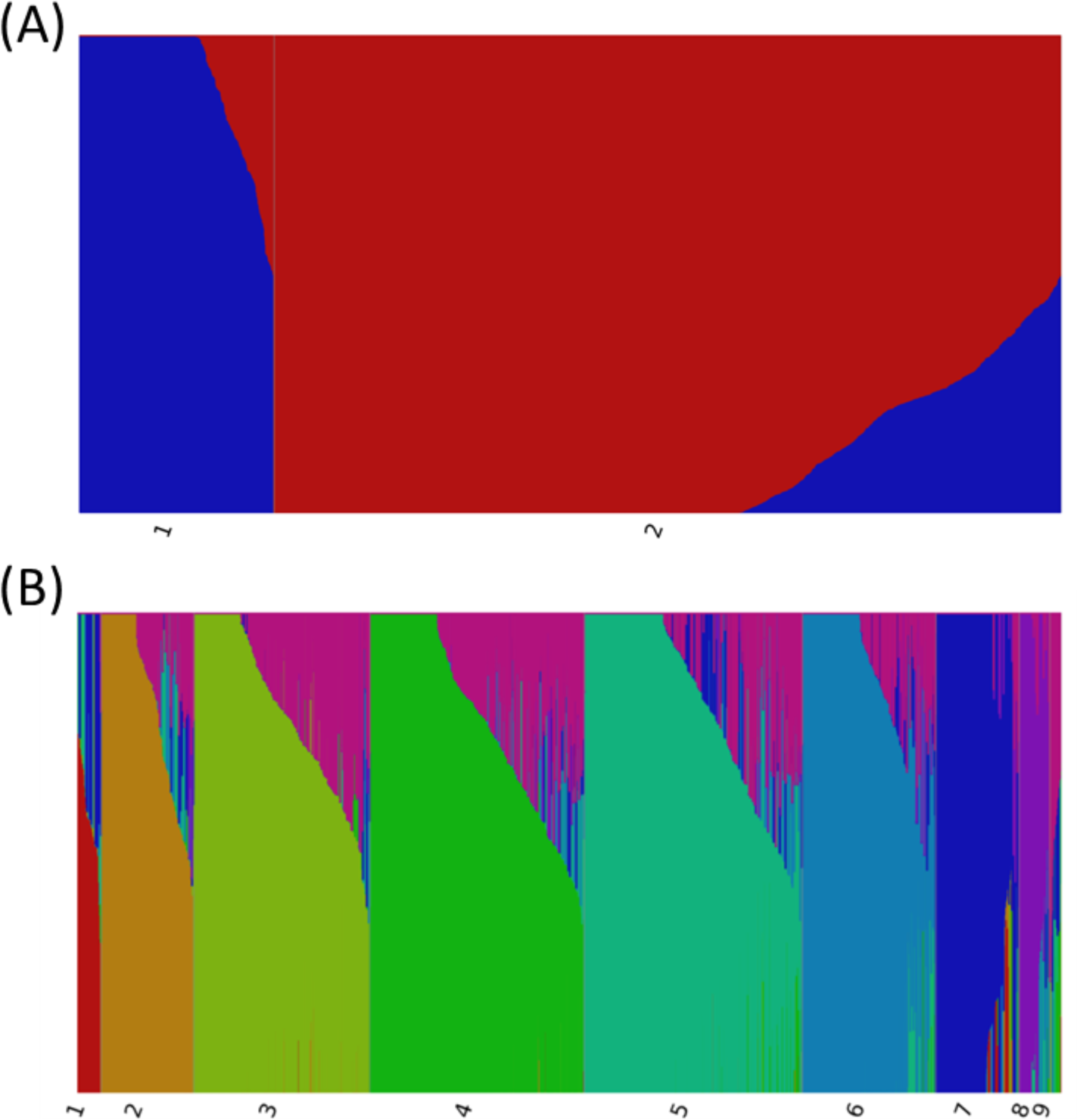
Population structure analysis of 2623 *B. napus* NAM panel based on 5743 SNP and SNaP markers. Population structure was shown as optimal model components (K) K=2 (A) and highest model components K=9 (B).

**Figure S4:**
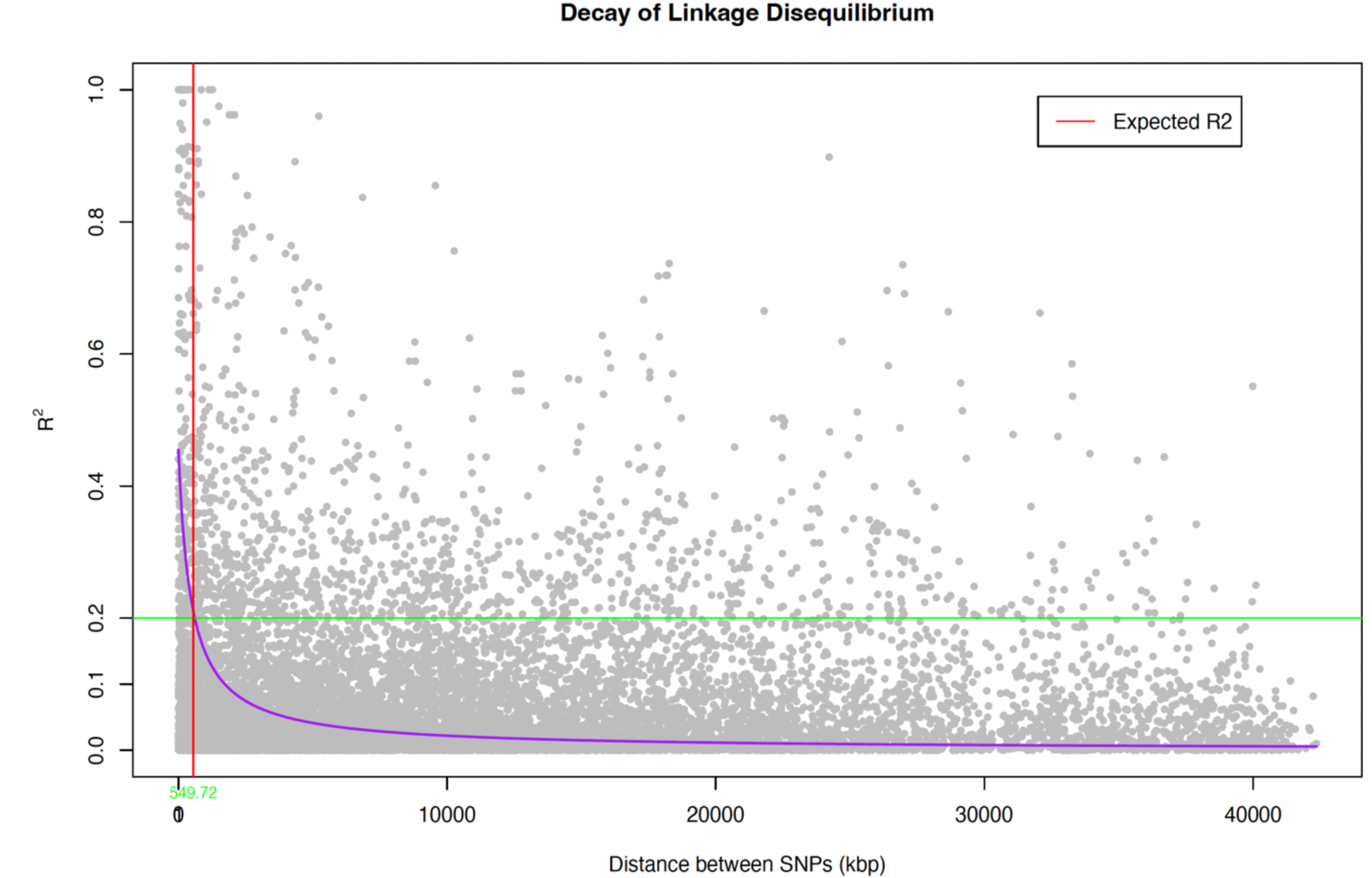
Linkage Disequilibrium (LD) decay plot with R^2^ values averaged within 10 Kb windows. The LD decay rate measured by the average pairwise correlation coefficient (*r*^2^) drops to the background LD (*r*^2^ = 0.21) ∼549.72 kb.

**Figure S5.**
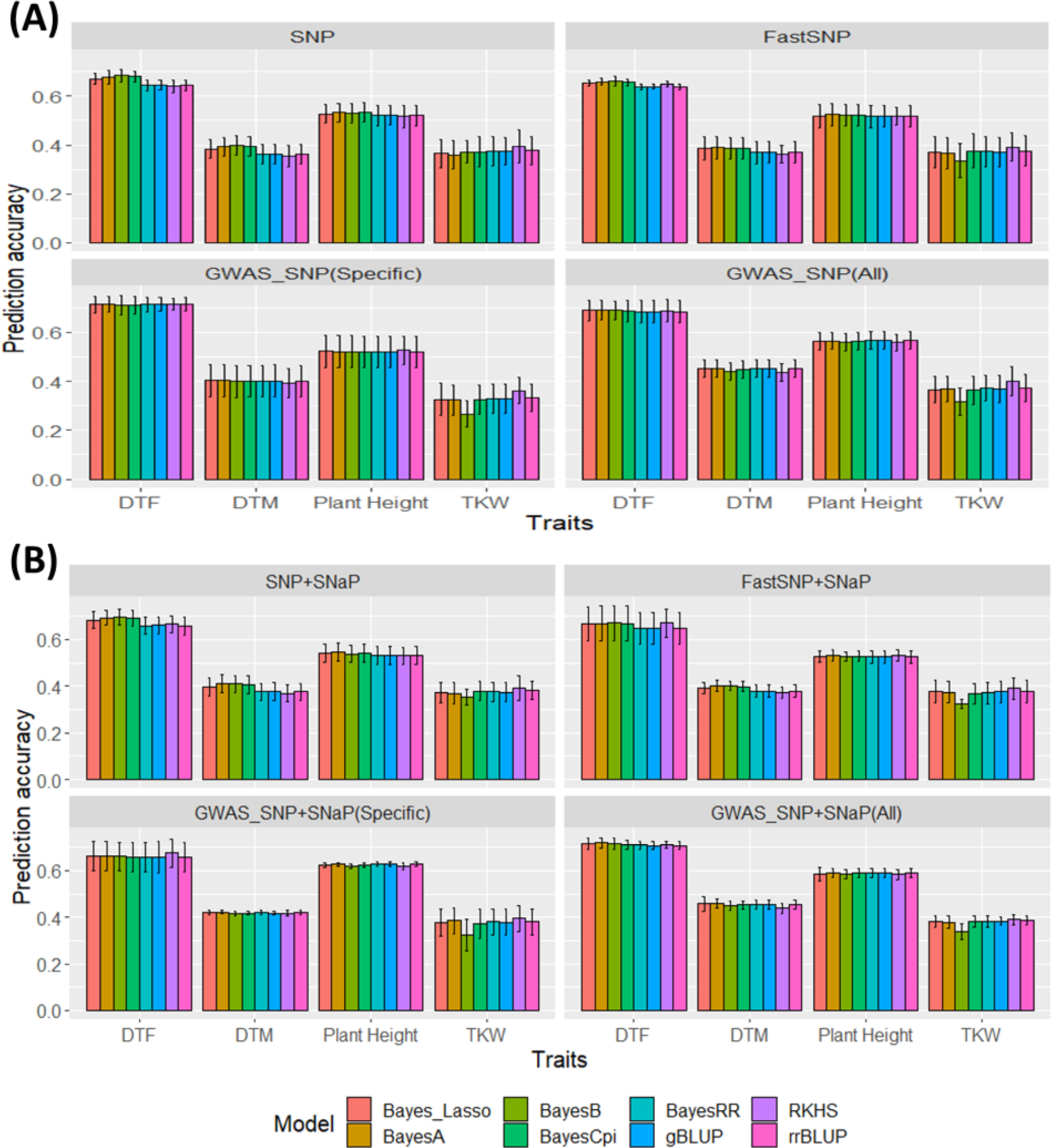
Genome prediction accuracy of *B. napus* NAM panel on 8 different marker sets using 8 different models. Prediction accuracy was estimated on marker sets developed from SNP markers (A) and combined SNP + SNaP markers (B).

**Figure S6:**
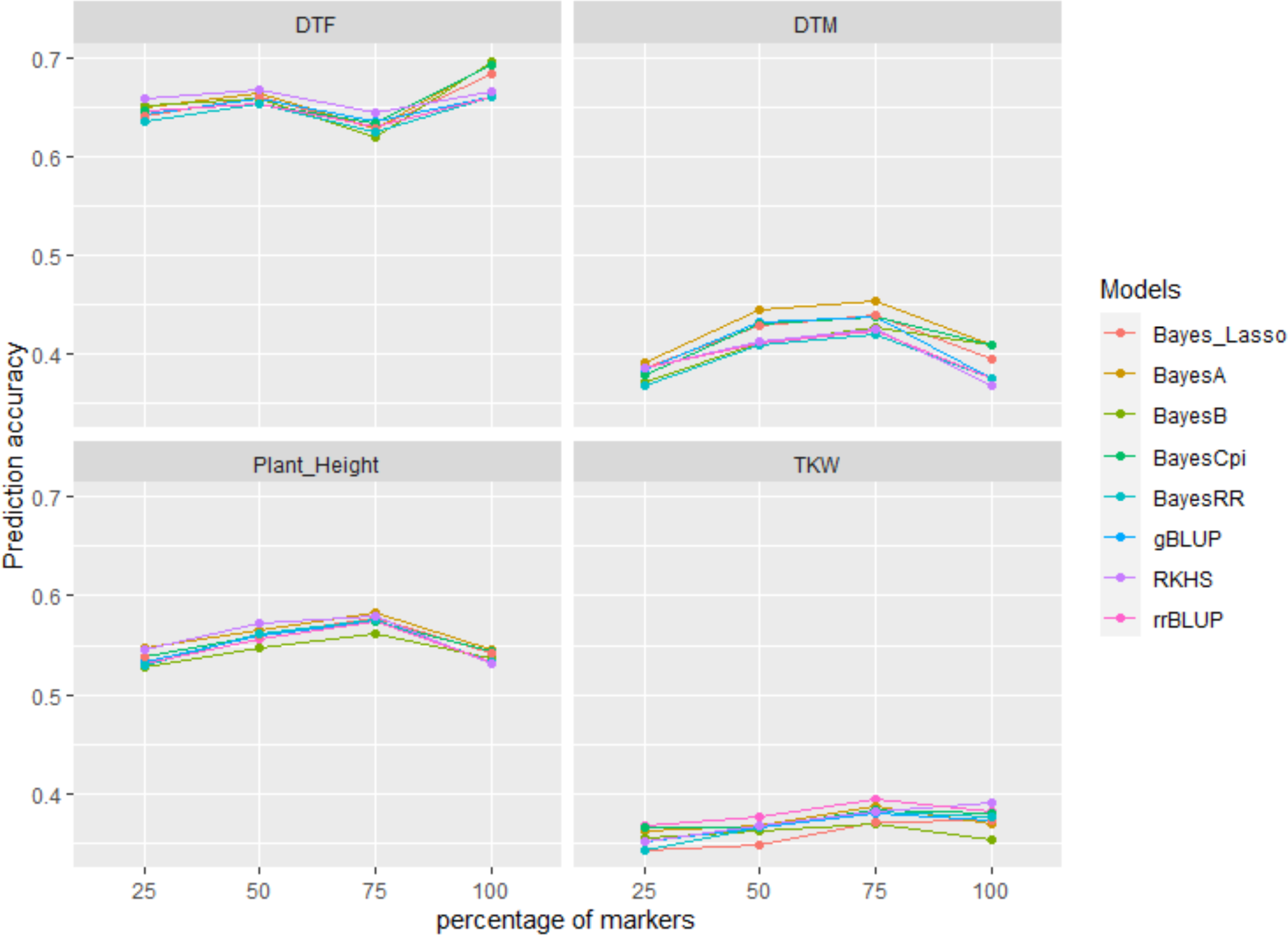
Maker density on prediction accuracy. All SNP+SNaP markers were evaluated on four different combinations (25% -100%) to estimate the genomic prediction accuracy of *B. napus* NAM panel.

**Figure S7:**
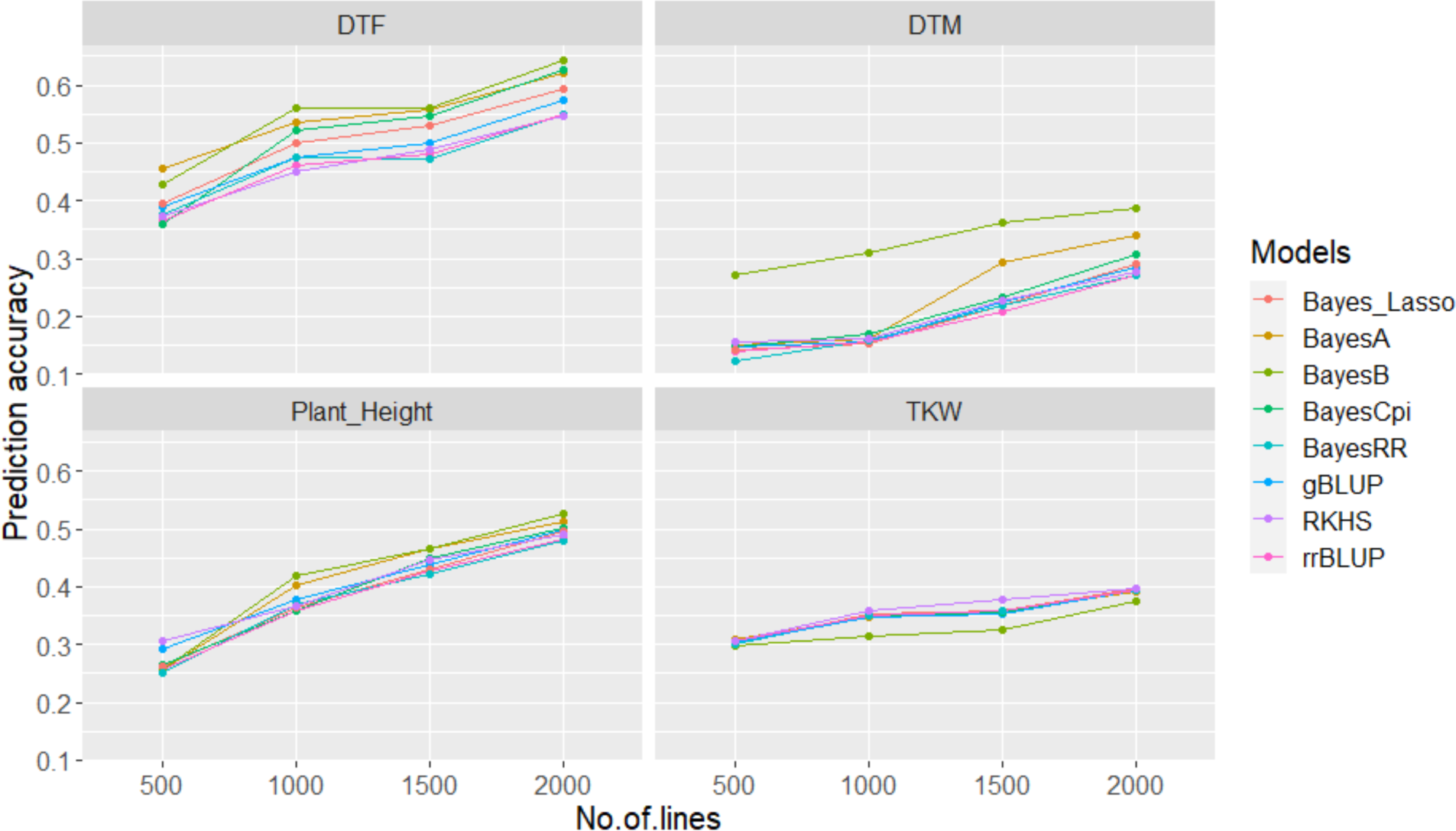
Training population size on prediction accuracy based on all SNP+SNaP marker set (5743 markers). Different set of lines (500, 1000, 1500 and 2000) were used to estimate the prediction accuracy on 500 lines using 8 different models.

## DATA AVAILABILITY STATEMENT

All datasets presented in this study are included in the article/Supplementary Material.

